# Recombinant Paraprobiotics as a New Paradigm for Treating Gastrointestinal Nematode Parasites of Humans

**DOI:** 10.1101/2020.09.02.278341

**Authors:** Hanchen Li, Ambily Abraham, David Gazzola, Yan Hu, Gillian Beamer, Kelly Flanagan, Ernesto Soto, Florentina Rus, Zeynep Mirza, Austin Draper, Sridhar Vakalapudi, Cheryl Stockman, Perry Bain, Joseph F. Urban, Gary R. Ostroff, Raffi V. Aroian

## Abstract

Gastrointestinal nematodes (GINs) of humans, *e.g*., hookworms, negatively impact childhood growth, cognition, nutrition, educational attainment, income, productivity, and pregnancy. Hundreds of millions of people are targeted with mass drug administration (MDA) of donated benzimidazole (BZ) anthelmintics. However, BZ efficacy against GINs is suboptimal, and reduced/low efficacy has been seen. Developing an anthelmintic for human MDA is daunting: it must be safe, effective, inexpensive, stable without a cold chain, and massively scalable. *Bacillus thuringiensis* (Bt) crystal protein 5B (Cry5B) has anthelmintic properties that could fill this void. Here we develop an API (Active Pharmaceutical Ingredient) form of Bt Cry5B compatible with MDA. We expressed Cry5B in asporogenous Bt during vegetative phase, forming cytosolic crystals. These **Ba**cteria with **C**ytosolic **C**rystals (BaCC) were rendered inviable (inactivated BaCC or IBaCC) with food-grade essential oils. IBaCC potency was validated *in vitro* against nematodes. IBaCC was also potent *in vivo* against human hookworm infections in hamsters. IBaCC production was successfully scaled to 350 liters at a contract manufacturing facility. A simple fit-for-purpose formulation to protect against stomach digestion and powdered IBaCC were successfully made and used against GINS in hamsters and mice. A pilot histopathology study and blood chemistry workup showed that five daily consecutive doses of 200 mg/kg Cry5B IBaCC (the curative single dose is 40 mg/kg) was non-toxic and completely safe. IBaCC is a safe, inexpensive, highly effective, easy-to-manufacture, and scalable anthelmintic that is practical for MDA and represents a new paradigm for treating human GINs.

## INTRODUCTION

Among the neglected tropical diseases, soil-transmitted helminths/nematodes (STHs/STNs) or gastrointestinal nematodes (GINs) collectively affect the largest number of people with a global estimate of >1.5 billion infected individuals (1). Human GINs include hookworms (*Ancylostoma duodenale, Ancylostoma ceylanicum, Necator americanus*), ascarids (*Ascaris lumbricoides*), and whipworms (*Trichuris trichiura) (2*). Human GIN parasites have an enormous impact on children, leading to physical growth stunting, cognitive impairment, malnutrition, anemia, impaired physical fitness, loss of future income, decreased educational attainment, and defective immune responses to other infectious diseases (*e.g*., HIV, malaria, tuberculosis) and vaccines (1–3). These parasites also cause significant complications for pregnant women and significant reductions in adult worker productivity, accounting for > 5 million disability-adjusted life years (DALYs) and productivity losses of over US$100 billion annually, with the majority of morbidity attributed to hookworms (2, 4–7).

Only one class of drug, the benzimidazoles (BZs), is approved and suitable for single-dose mass drug administration (MDA) (8, 9). GIN resistance to these drugs develops readily and is extremely common in veterinary medicine, where they have been used much longer and more intensely than in human medicine (8, 10). Against human GINs, these drugs have poor efficacy against whipworms. Low efficacy of albendazole (the most efficacious BZ used in humans) against hookworms and *Ascaris* has been reported in many locales, with definitive BZ resistance alleles detected in natural populations of human hookworms in Kenya and Brazil (11–20).

New mechanism-of-action anthelmintics are urgently needed in the pipeline as the lead time for drugs to reach the market is years. As human GIN MDA intensifies, further loss of BZ efficacy is highly likely. However, developing new drugs for humans GINs is exceedingly difficult: 1) drug development is very expensive, costing several billions (21), a cost that cannot easily be recouped on diseases of the poorest peoples; 2) the number of people impacted is enormous -- any new therapy has to be massively scalable (an estimated 1.5 billion doses currently needed for children and women alone) and cheap (BZs are currently donated) (22); 3) the therapy has to withstand harsh environmental conditions without a cold chain; and 4) the therapy has to be safe and effective. Thus, the normal rules of “market-incentive” drug development do not apply. In fact, no drug has ever been developed for human GINs; all drugs used are expanded label uses of drugs developed for veterinary medicine. Of the two drugs used in MDA today, albendazole was approved in humans in 1982 (23); mebendazole in 1974 (USA; (24)). No new drugs have thus entered human GIN treatment for more than 30 years.

*Bacillus thuringiensis* (Bt) crystal (Cry) proteins have the potential to offer a novel, natural, safe, and broad spectrum anthelmintic alternative. Bt spores are mass produced globally as a biopesticide, encompassing ~75% of the bioinsecticide market (25). The main insecticidal components of Bt are three-domain (3D-)Cry proteins that bind specifically to the invertebrate intestine, damaging the gut and killing the invertebrate target (26). 3D-Cry proteins have also been engineered into a range of food crops and are expressed in ~100 mHa of transgenic crops worldwide (27). More than a dozen different 3D-Cry proteins have been tested and found to be completely safe to vertebrates at doses of >>1000 mg/kg, and are FDA/EPA-approved for ingestion (28, 29). The 3D-Cry protein Cry5B is related to the 3D-Cry proteins used as insecticides but targets nematodes instead., When administered orally as spore-crystal lysates or SCLs (a mix of Bt spores and Cry protein crystals as they naturally occur), Cry5B is highly effective against GIN infections in hamsters, pigs, and dogs (30–34). The nematode receptor for Cry5B is an invertebrate-specific glycolipid absent in vertebrates (35).

Developing 3D-Cry proteins as large-scale ingested therapeutics compatible with human MDA, however, requires a very different set of considerations than application as topical or transgenic biopesticides and insecticides. Here we address these considerations, developing an active pharmaceutical ingredient (API) based on Cry5B compatible with safety, cost, scale, ease of production, and stability required for human MDA.

## RESULTS

### Inactivated Bacterium with Cytosolic Crystal (IBaCC)

We reasoned that delivering Cry5B in a live bacterium (Bt or otherwise) to humans would be problematical given that 1) Bt is closely related to *Bacillus cereus*, which can cause food poisoning, 2) release of live recombinant bacteria has environmental concerns because live bacteria in the soil could select for Cry5B resistance against free living stages of hookworms t, 3) live bacteria could replicate in the human gastrointestinal tract (amplifying environmental and resistance concerns), 4) there are uncertainties in the response of billions of people to live bacteria, 5) degradation of live bacteria during storage could reduce potency and stability, and 6) delivery of a stable live bacterial therapeutic around the world for MDA would be difficult to achieve. An alternative of producing and delivering Cry5B as a purified protein is also problematic as making enough purified protein cheaply and massively for MDA is difficult to envision.

We therefore set out to deliver Cry5B, without purification of the protein, as part of a dead bacterial product (or “paraprobiotic” (50)). To make a Cry5B paraprobiotic, we initially turned to and produced Cry5B spore-crystal lysates, which have been used for most published *in vivo* studies of Cry5B anthelmintic efficacy as well as all Bt insecticidal production and use (31, 33, 34, 43, 45). We attempted to inactivate (kill) the spores without losing activity of the crystal protein using gamma irradiation (51). Although gamma irradiation resulted in a significant (10^6^ - 10^7^) reduction in spore viability (Fig. 1A), the procedure completely killed CryB anti-nematode activity (Fig. 1B). Preliminary studies using chemical treatment instead of gamma-irradiation to inactive spores without damaging crystal activity yielded similar, disappointing results.

**Figure 1.**
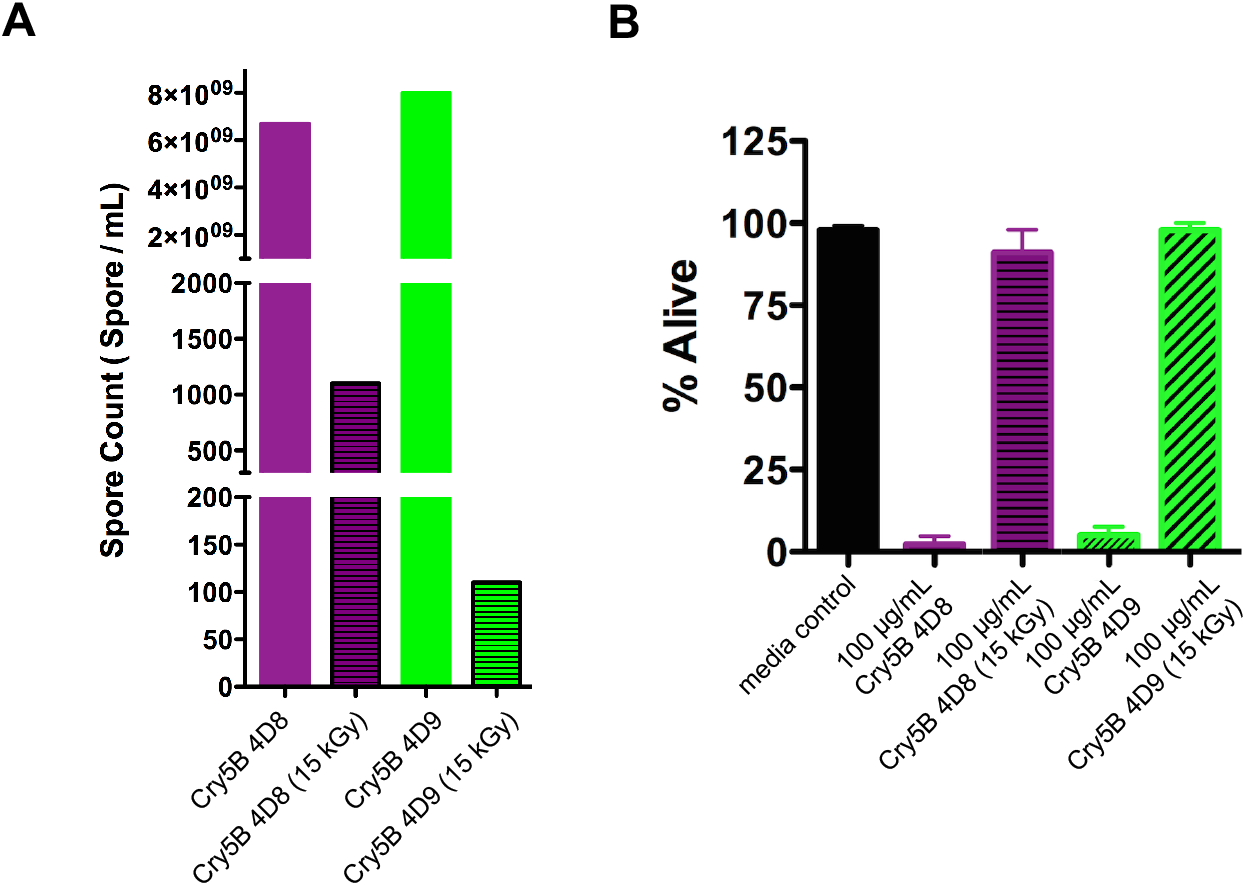
Effects of gamma irradiation on Bt spore viability and Cry5B bioactivity. (A) Effect of 15 kGy of gamma irradiation on spore counts for two different Cry-Bt strains (4.D.8, 4.D.9) transformed with a Cry5B expressing plasmid. (B) Comparison of Cry5B efficacy expressed in 4.D.8 and 4.D.9 on *C. elegans* viability at 100 μg/mL before and after 15 kGy irradiation. Labels without “(15 kGy)” indicate non-gamma irradiated samples. Labels with “(15 kGy)” indicate gamma-irradiated samples.

We therefore decided that a better approach would be to express Cry5B in vegetative bacteria that could be more easily killed (inactivated). Based on our previous experiences and based on the fact that the inactivation process needed to be compatible with human ingestion, we hypothesized that food-grade monoterpenes derived from essential oils would be effective and safe antimicrobials for this application (52–55). We therefore expressed Cry5B in asporogenous (spo0A-) Bt so that the Cry5B crystals would be formed inside of a vegetative Bt without subsequent spore formation (Fig. 2A) (46, 48). We call such cells BaCC for **Ba**cterium with **C**ytosolic **C**rystal. A number of food-grade monoterpenes were found that were capable of inactivation (killing) of spo0A-cells (Table S1). When Cry5B was expressed in these cells and inactivated with terpene, the crystals stayed trapped within the cells, and the protein remained intact (Fig. 2A, B). A >10^7^ fold reduction in CFUs was seen upon terpene treatment, with often no viable cells detected (Fig. 2C). We call these cells IBaCC for **I**nactivated BaCC.

**Figure 2.**
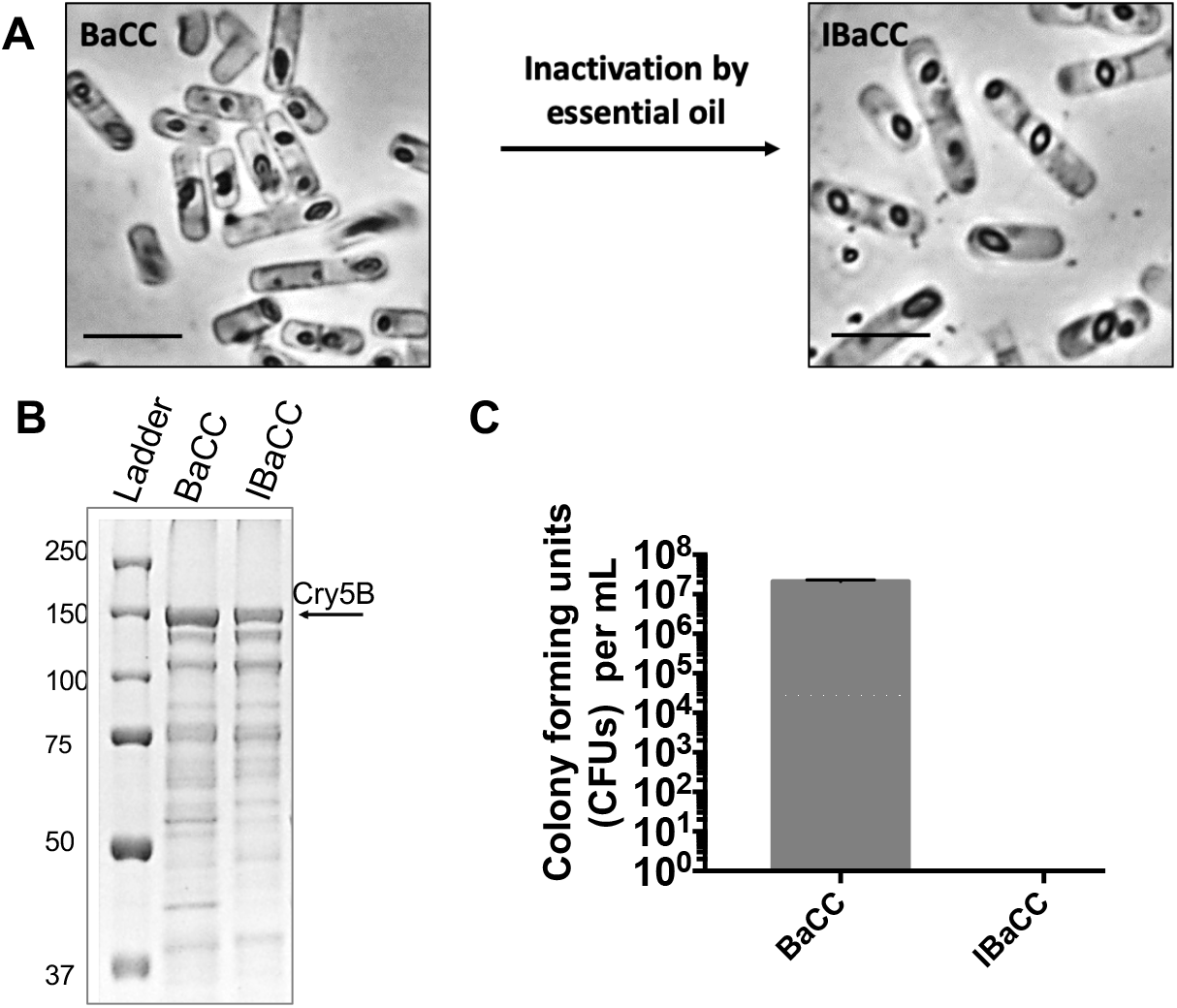
BaCC (Bacterium with cytosolic crystal) and IBaCC (Inactivated Bacterium with Cytosolic Crystal). (A) spo0A-Bt cells expressing Cry5B from a vegetative promoter before (BaCC) and after (IBaCC) treatment with essential oil. Cry5B bipyramidal crystals (dark) are evident inside the bacteria pre- and post-treatment. Scale bar = 5 μm. (B) Protein gel showing Cry5B protein expressed in spo0A-cells before and after essential oil treatment. (C) Spore counts from spo0A-cells expression Cry5B before and after essential oil treatment along with standard deviation (actual spore counts = 2.1 x 10^7^ CFU/mL in BaCC and 0 in IBaCC).

### IBaCC is an active nematicide

IBaCC was tested and quantified for anti-nematode activity initially against free-living stages of nematodes. Against the free-living nematode *C. elegans*, IBaCC (containing Cry5B crystals) intoxicated and killed L4/adult stages whereas identically prepared IBa (**I**nactivated **Ba**cterium with empty vector control; no Cry5B) did not (Fig. 3A). When tested against the free-living developing larval stages of the human hookworms *A. ceylanicum* and *N. americanus*, Cry5B IBaCC, but not vector-only IBa, was highly potent, strongly inhibiting larval development even at doses of 0.5 - 1 μg/mL (Fig. 3B). We next tested IBaCC for anti-nematode activity against adult hookworm parasites *in vitro*. Cry5B IBaCC, but not IBa, was potent at intoxicating both species of adult parasitic hookworms at doses as low as 5 μg/mL (Fig. 4A, B). Cry5B in IBaCC showed a dose-dependent inhibition of motility similar to previously published studies with purified Cry5B (30). We had previously confirmed uptake of 0.4 μm particles by adult hookworms (56). To confirm uptake of IBaCC by hookworms, we labeled IBaCC with rhodamine, which predominantly labels full length Cry5B. (Fig. 4C; rhodamine labeled IBaCC was fully potent as seen by 100% dead adult hookworms after 24 hours in 16 μg/mL Cry5B rhodamine-IBaCC). Visualization of uptake of rhodamine IBaCC after 4 hours by adult hookworms *in vitro* was confirmed by fluorescence microscopy (Fig. 4D). Taken together, these data indicate that Cry5B expressed in IBaCC is ingested by, and is highly active against, nematodes, even though the bacterium is not viable. Conversely, empty vector inactivated bacteria (IBa) without Cry5B is not active against nematodes.

**Figure 3.**
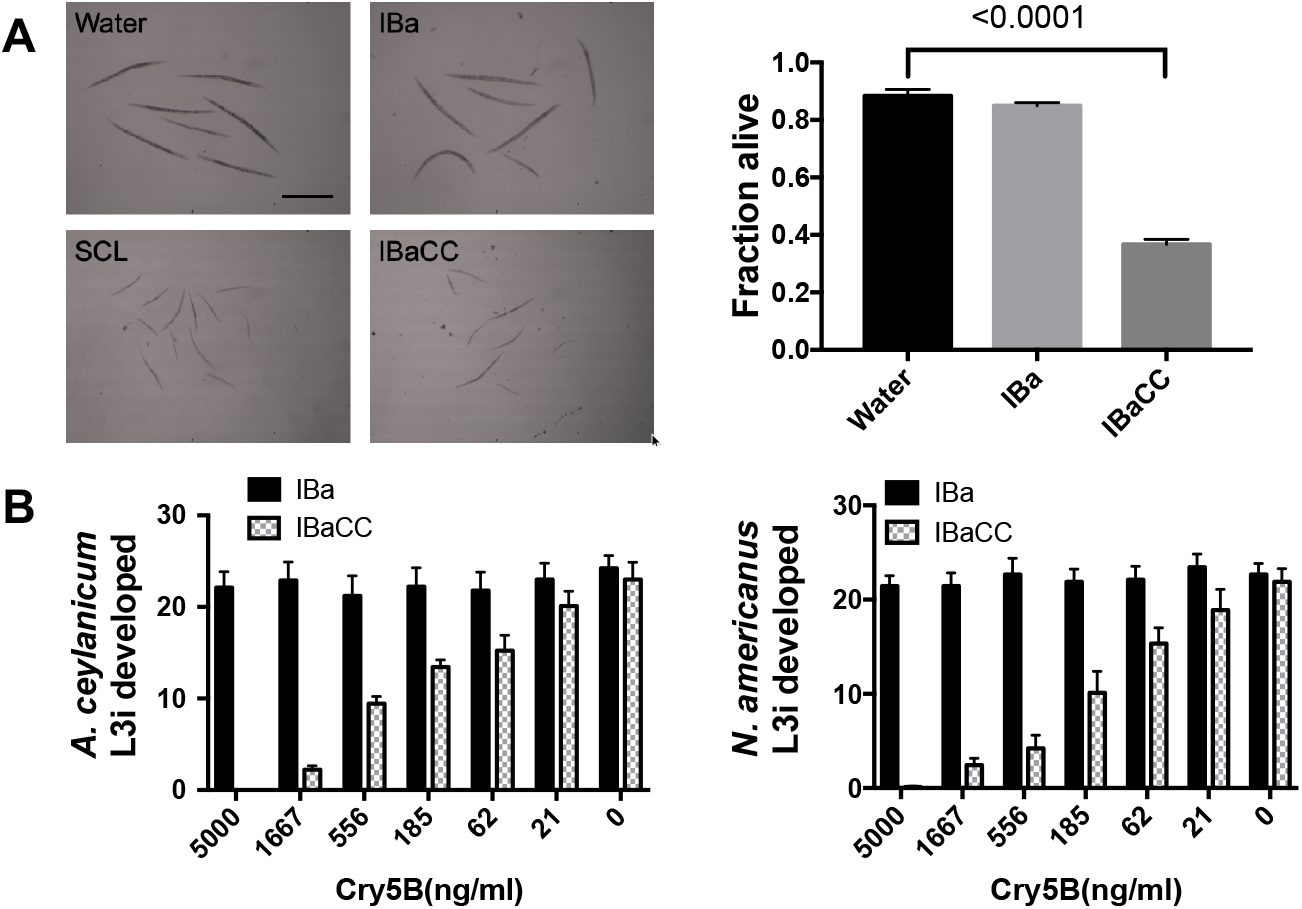
Efficacy of IBaCC *in vitro* against nematodes. (A) *C. elegans*. Left, photos of *C. elegans* N2 exposed to various conditions (spore crystal lysate or SCL and IBaCC; both = 40 μg/mL Cry5B). Scale bar = 200 μM. The *C. elegans* were treated with azide to immobilize them just prior to imaging. Right, viability of *C. elegans glp-4(bn2)* L4 hermaphrodites under various conditions (IBaCC = 29 μg/mL Cry5B). P value for comparison is one-tailed T-test. (B) Hookworm larval development. Plotted are the numbers of L3i larvae that developed from 60 hookworm eggs within 7 days (*A. ceylanicum*, left; *N. americanus*, right). X axis indicates concentration of Cry5B for IBaCC (IBa = 0 for all).

**Figure 4.**
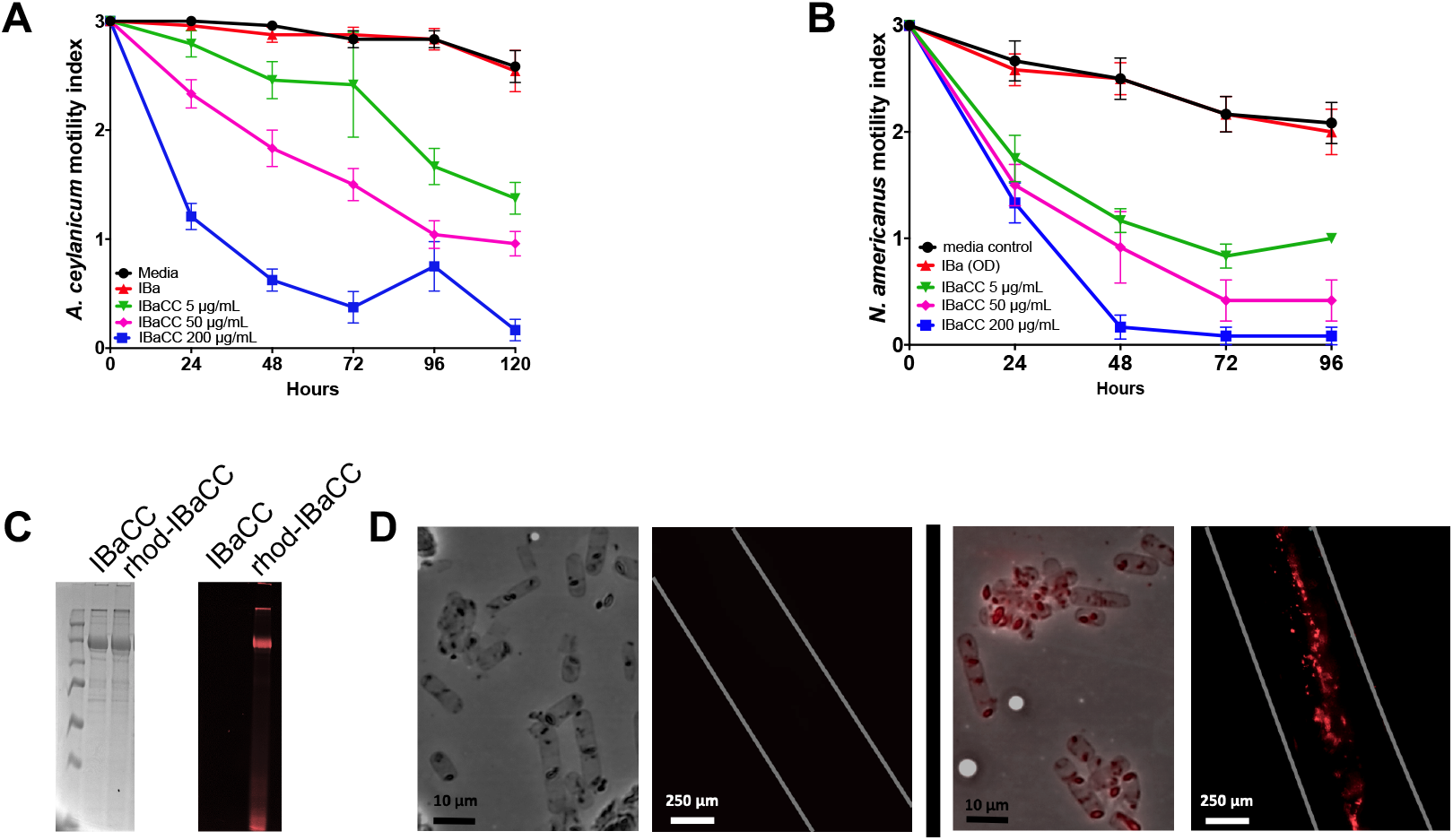
Impact of Cry5B IBaCC on adult hookworms *in vitro*. Adult hookworm motility over time at 5, 50, and 200 μg/mL Cry5B for (A) *A. ceylanicum* adults and (B) *N. americanus* adults (right) *in vitro* averaged over three independent trials. In all figures, plots show average and standard error. (C) left: SDS PAGE showing IBaCC before (IBaCC) and after (rhod-IBaCC) labeling with rhodamine; right: UV fluorescent image of same gel, showing predominant labeling of full length Cry5B band in rhod-IBaCC. (D) Uptake of rhod-IBaCC. Left pair of images, bright field image of IBaCC and rhodamine fluorescent image of adult hookworm fed IBaCC for 4 hours; Right pair of images, bright field image of rhod-IBaCC and rhodamine fluorescent image of adult hookworm fed rhod-IBaCC for 4 hours. Uptake of rhodamine-labeled Cry5B crystals is evident.

### Cry5B IBaCC is a potent anthelmintic *in vivo* against both genera of human hookworms

We next tested whether or not IBaCC efficacy *in vivo* against hookworms. *Ancylostoma ceylanicum* is an important zoonotic hookworm parasite of humans (57–60), and *A. ceylanicum* infections in hamsters are considered a good laboratory model for hookworm infections in humans (61). *A. ceylanicum* is also in the same genera as *Ancylostoma duodenale*, the second most common hookworm parasite in humans. Hamsters were infected with *A. ceylanicum*, and the infestations were allowed to proceed to mature adults with the appearance of parasite eggs (fecundity) excreted into the hamster feces (31, 33, 44). These hamsters were then treated with IBaCC produced in our laboratory containing 10 mg/kg Cry5B (Figure 5A) or treated with IBa (vector-only control produced at the same time and processed identically to Cry5B IBaCC but lacking Cry5B). Whereas IBa had no impact on hookworm burdens or fecal egg counts (Fig. 5A, Supplemental Fig. 1; Table 1), IBaCC had a strong impact, resulting in elimination of more than 93% of the hookworm (Fig. 5A; Table 1).

**Figure 5.**
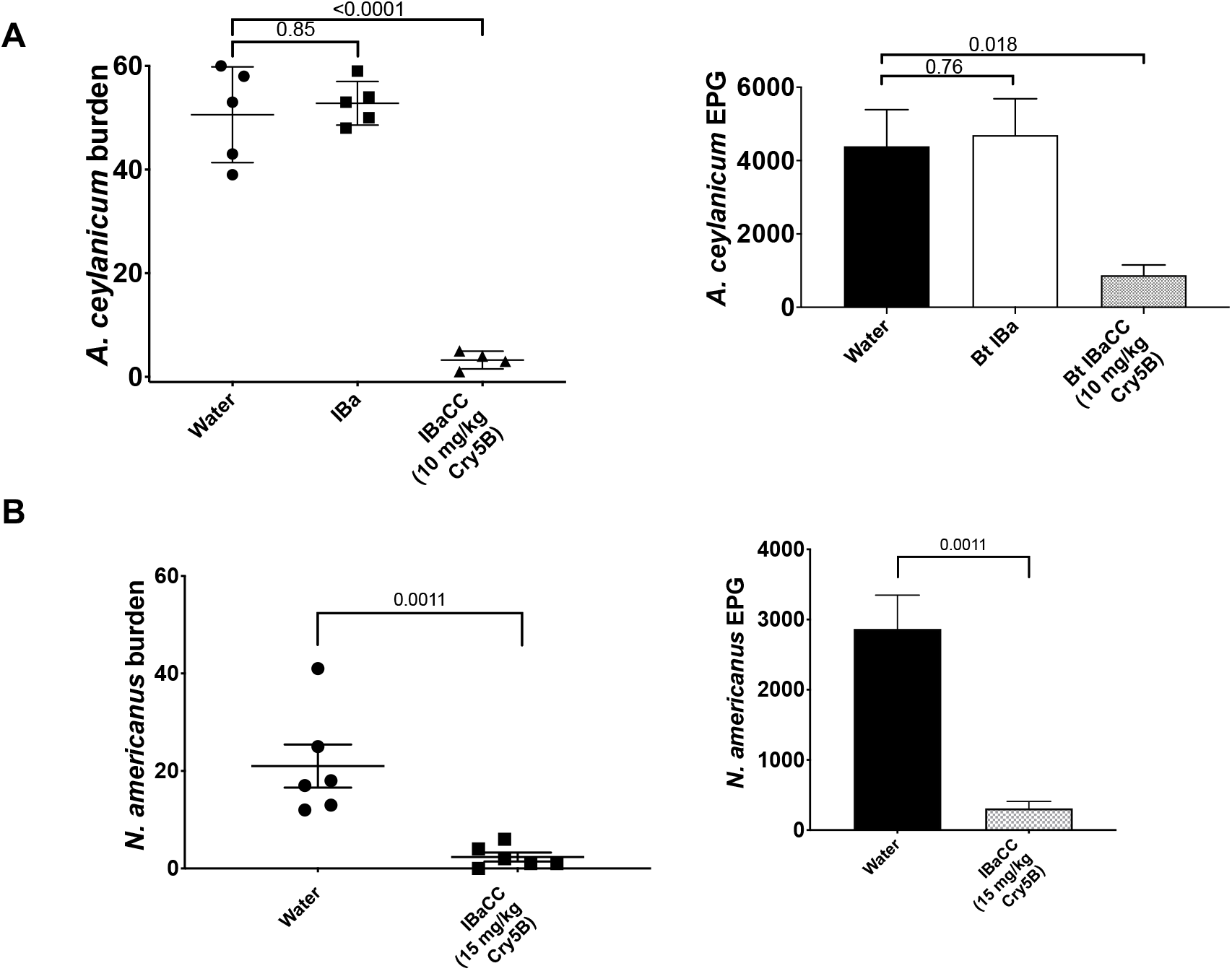
Efficacy of IBaCC produced in the laboratory *in vivo* against both human hookworm genera. (A) Shown are mean *A. ceylanicum* hookworm burdens (left) and fecal egg counts (right) in infected hamsters treated with water control, IBa, or IBaCC containing Cry5B. Here and subsequent figures error bars are standard error. EPG = eggs per gram of feces. (B) Mean *N. americanus* hookworm burdens (left) and fecal egg counts (right) in infected hamsters treated with water control or IBaCC containing Cry5B.

**Table 1.** Data from *in vivo* experiments.

| Figure | Condition | Average worm burden (n) | Change from control | Average EPG (n) | Change from control |
| --- | --- | --- | --- | --- | --- |
| 5A | Water | 50.6 (5) | NA | 4390 (5) | NA |
|  | IBa | 52.8 (5) | 4.30% | 4695 (5) | 6.90% |
|  | IBaCC 10 mg/kg Cry5B | 3.3 (4) | -93.50% | 875.0 (4) | -80.10% |
| 5B | Water | 21.0 (6) | NA | 2867 (6) | NA |
|  | IBaCC 15 mg/kg Cry5B | 2.3 (6) | -89.00% | 308.3 (6) | -89.20% |
| 6A | Water | 44.3 (6) | NA | 2275 (6) | NA |
|  | IBaCC 2 mg/kg Cry5B | 14.5 (6) | -67.30% | 808.3 (6) | -64.50% |
|  | IBaCC 6 mg/kg Cry5B | 5.2 (6) | -88.30% | 366.7 (6) | -83.80% |
| 6B | Water | 35.6 (5) | NA | 2790 (5) | NA |
|  | IBaCC 15 mg/kg Cry5B | 0.4 (5) | -98.90% | 30.0 (5) | -98.90% |
| 7A | Water | 21.8 (5) | NA | 3210 (5) | NA |
|  | IBaCC 2.4 mg/kg Cry5B | 8.4 (5) | -61.50% | 1220 (5) | -62.00% |
|  | IBaCC 2.4 mg/kg Cry5B + Bicarb | 4.2 (5) | -80.70% | 850.0 (5) | -73.50% |
| 7B | Water | 42.3 (6) | NA | 3146 (6) | NA |
|  | IBaCC 6 mg/kg Cry5B | 3.8 (6) | -91.00% | 404.2 (6) | -87.20% |
|  | FD-IBaCC 6 mg/kg Cry5B | 1.0 (6) | -97.60% | 241.7 (6) | -92.30% |
| 8 | Water | 60.2 (6) | NA | 6517 (6) | NA |
|  | Bicarb | 46.5 (6) | -22.80% | 8325 (6) | 27.70% |
|  | FD-IBaCC 40 mg/kg Cry5B + Bicarb | 28.3 (6) | -53.00% | 883.3 (6) | -86.40% |
| Supplement | Water | 14.7 (6) | NA |  |  |
|  | Bicarb only | 17.0 (6) | 15.60% |  |  |
|  | IBa | 16.7 (6) | 13.60% |  |  |

We also tested Cry5B IBaCC against the most common hookworms of humans, *N. americanus*, which can be maintained and studied in immunosuppressed hamsters (31). This hookworm infection is more difficult to treat (*e.g., Necator*, but not *Ancylostoma* hookworms, are recalcitrant to ivermectin treatment; (62)). Cry5B IBaCC was highly effective against *N. americanus* hookworm infections in hamsters (Fig. 5B; Table 1).

### IBaCC is scalable and transferrable

We then looked at whether Bt fermentation and Cry5B production and processing to IBaCC could successfully be transferred to and scaled up at a contract manufacturing facility. Our IBaCC strain and protocols were transferred to a contract manufacturing organization (CMO) at Utah State University. Fermentation of Cry5B BaCC and processing to IBaCC was brought up to the 350 L scale by CMO. Cry5B IBaCC produced at the CMO was then tested *in vivo* against *A. ceylanicum* hookworm infestations in hamsters at 2 and 6 mg/kg Cry5B. IBaCC produced at the manufacturing facility was effective at reducing *A. ceylanicum* burdens and parasite fecal egg counts in hamsters relative to water control (Fig. 6A; Table 1). Increasing the dose of Cry5B in this large-scale IBaCC production run to 15 mg/kg essentially cured the parasite infestation (Fig. 6B; Table 1). Efficacy was similar to that achieved with in-house produced IBaCC (Fig. 5 compared to Fig. 6).

**Figure 6.**
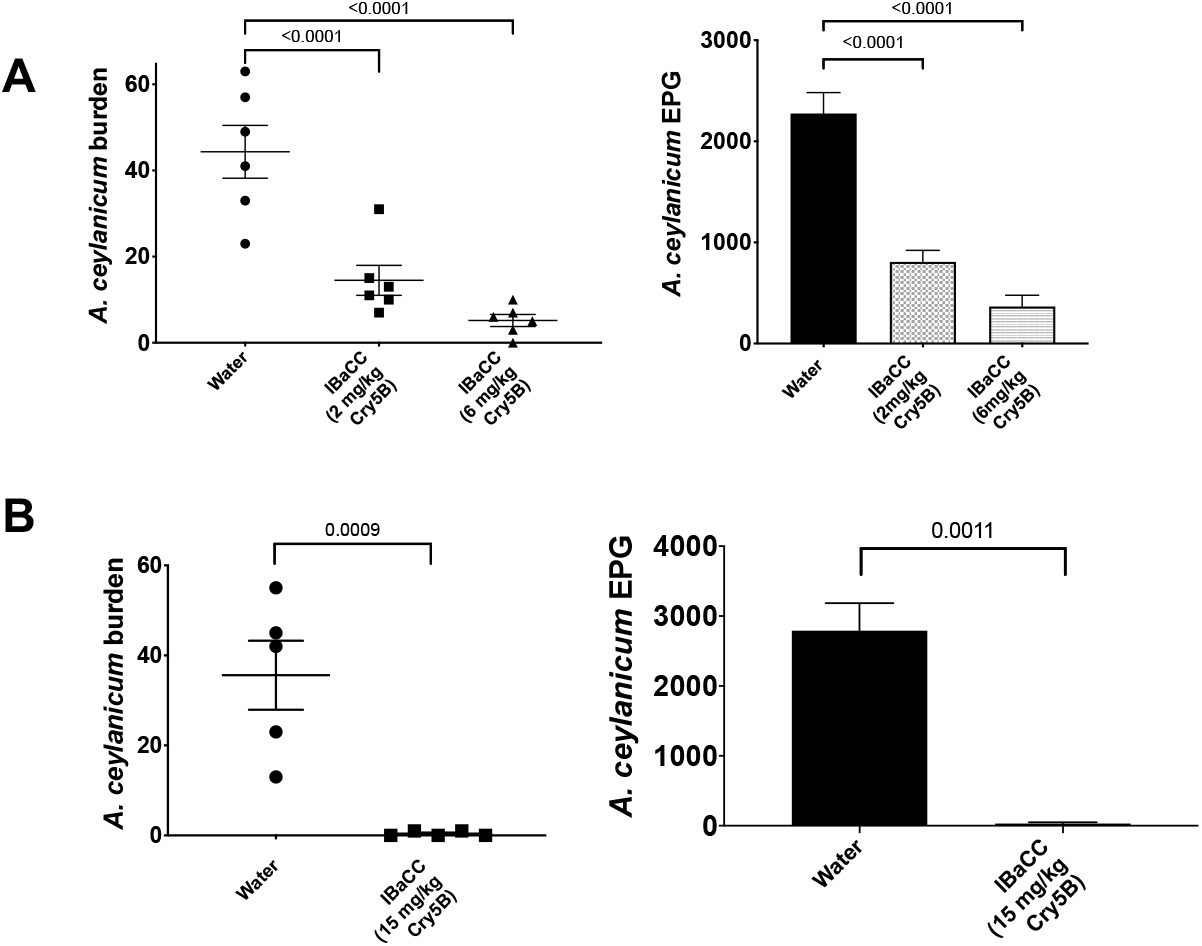
Efficacy of Production Facility IBaCC *in vivo* against *A. ceylanicum* hookworms. (A) Dose response of *A. ceylanicum* burdens (left) and fecal egg counts (right) in infected hamsters treated with water control or CMO-produced IBaCC containing Cry5B. (B) *A. ceylanicum* burdens (left) and fecal egg counts in infected hamsters treated with water or CMO-produced 15 mg/kg Cry5B in IBaCC. EPG = eggs per gram of feces. P values for relevant comparisons are given.

### Cry5B IBaCC can be made into a fit-for-purpose formulation and dried down

Previous work has shown that the potency of Cry5B spore crystal lysates are enhanced by addition of a pre-treament with cimetidine to neutralize stomach acid. However, pre-treatment with a drug like cimetidine is not compatible with MDA. We therefore tested whether the development of a safe and simple “fit-for-purpose” formulation compatible with MDA, notably simultaneous delivery with sodium bicarbonate, would be protective for IBaCC. IBaCC was given alone or simultaneously with sodium bicarbonate to hookworm-infected hamsters. We found that acid neutralization with sodium bicarbonate given simultaneously with IBaCC slightly but significantly increased IBaCC efficacy against hookworms (Fig. 7A; Table 1).

**Figure 7.**
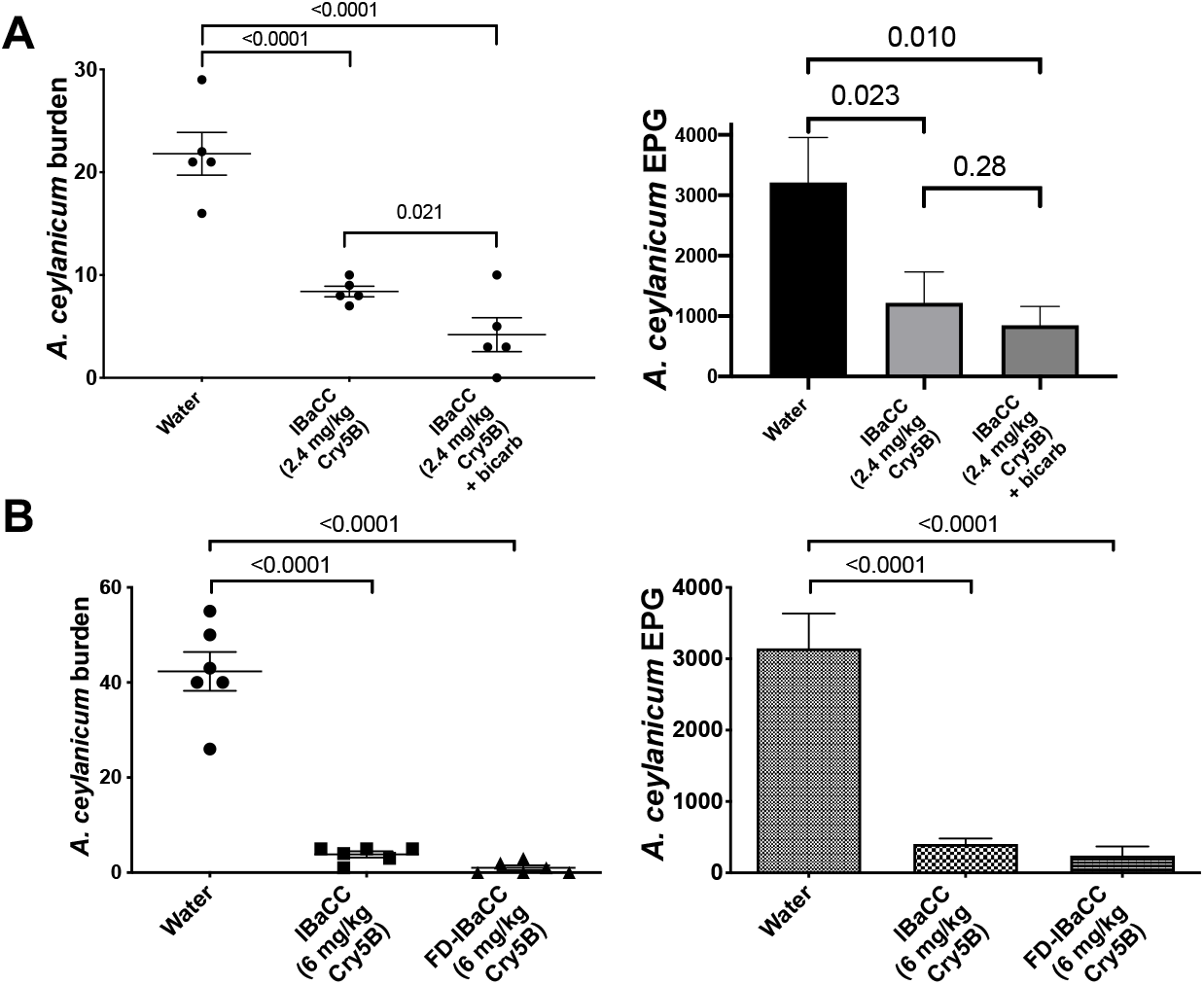
Fit-for-purpose formulation studies for Cry5B IBaCC. (A) Mean *A. ceylanicum* hookworm burdens (left) and fecal egg counts (right) in infected hamsters treated with water control, IBaCC, or IBaCC mixed with sodium bicarbonate. No cimetidine pre-gavage was used in these experiments. (B) Mean *A. ceylanicum* hookworm burdens (left) and fecal egg counts (right) in infected hamsters treated with water control, IBaCC, or the same batch of IBaCC freeze-dried (FD). P values for relevant comparisons are given. One-tailed Dunnett’s; one-tailed T test.

Delivery of IBaCC as a powder (and not as a liquid slurry as in above experiments) is also a critical parameter for storage and MDA. We therefore freeze-dried IBaCC into a powder and compared the efficacy of the same batch of IBaCC before and after freeze-drying (given *per os* as a powder suspension in water). As shown (Fig. 7B; Table 1), freeze-drying had no impact on IBaCC *in vivo* efficacy.

Efficacy of freeze-dried IBaCC simultaneously delivered with sodium bicarbonate was also tested against a different, luminal-feeding (63) intestinal parasitic nematode in a different host, namely *Heligmosomoides polygyrus bakeri* infections in mice. These data confirm that this simple fit-for-purpose IBaCC formulation is effective against a different parasitic nematode in a number of different host (Fig. 8). The single dose efficacy seen (53% reduction single dose 40 mg/kg Cry5B) was superior to that shown in previous studies with this parasite (31, 45).

**Figure 8.**
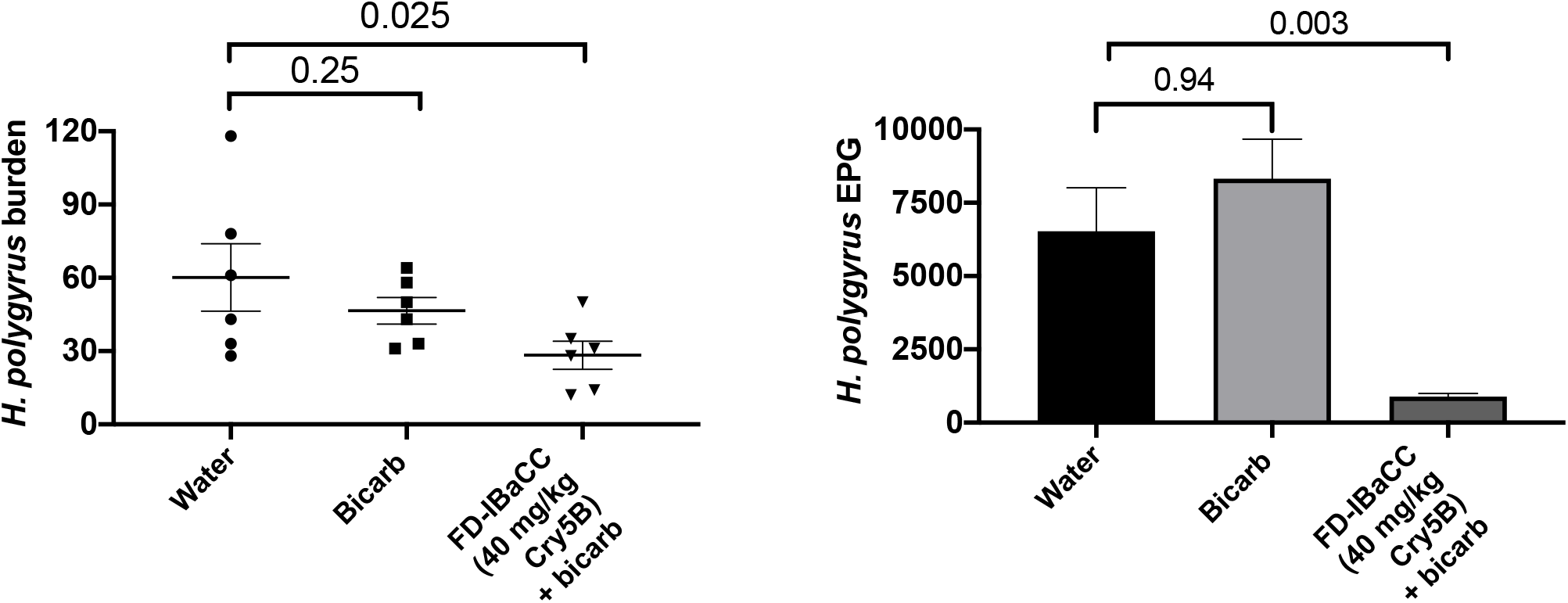
Confirmation of fit-for-purpose Cry5B IBaCC efficacy against a different, luminal feeding parasite in a second host. Mean *H. polygyrus bakeri* burdens (left) and fecal egg counts (right) in infected mice treated with water, sodium bicarbonate, or single-dose freeze-dried (FD) Cry5B IBaCC mixed with sodium bicarbonate. The FD-IBaCC used was an independent batch from that in Figure 7.

### Pilot preclinical toxicology study

Cry proteins have a stellar safety record and are considered non-toxic even at high doses, and the bacterium delivering IBaCC is dead (inactivated). Thus, IBaCC is predicted to be completely safe. However, IBaCC represents a new form for delivering a Cry protein. Thus, we performed a pilot maximal Cry5B dose safety study (Fig. S2) using histopathology blood chemistry as readouts.

Sixteen uninfected hamsters (eight females and eight male) were split into two groups. Half the hamsters (four females and four males) were given 200 mg/kg Cry5B IBaCC *per os* daily for five days in a row. Since a single 40 mg/kg dose of Cry5B is curative for hookworms (31), each dose represents 5X the curative dose and, cumulatively, 5X the required number of doses were given. The other half of the hamsters received an equal volume of water *per os* on each of the five days. Three days after the final treatment, half of the hamsters in each group were sacrificed (acute group). All major organs were immediately dissected, fixed in formalin, sectioned, and stained, looking for signs of disease and lesions (see Methods for details). Ten days after the final treatment, the remaining half of the hamsters in each group were sacrificed (recovery group). The acute group would permit observation of any short-term adverse consequences of treatment, whereas the recovery group would permit observation of the resolution of adverse consequences seen in the acute group (if any). After staining, 274 sections from all major organs and tissues were examined blinded and scored by a board-certified pathologist.

The full results are presented in Table S2. Based on this maximal dose pilot treatment study, there were no significant differences seen between water and IBaCC groups. There were also no significant differences seen between the acute and recovery groups comparing across other groups, and no significant differences seen between males and females comparing across other groups. No significant pathologies were seen in any groups.

After sacrifice, blood samples were also collected and analyzed for blood chemistry with a focus on enzymes that could be indicative renal or hepatic injury (alanine aminotransferase or ALT, gamma-glutamyl transferase or GGT, aspartate aminotransferase or AST, bilirubin, blood urea nitrogen or urea, and creatinine).

Comparison of hamsters in the water versus Cry5B IBaCC groups showed no statistical difference in any of these levels (Table S3), consistent with high level of safety and lack of toxicity. Repeated, maximal dosing of IBaCC appear to be completely safe and non-toxic based on this pilot histopathology and blood chemistry study. Acute and chronic GLP toxicology studies with larger sample sizes are planned in the future with clinical-grade Cry5B IBaCC.

## DISCUSSION

The primary aim of this study was to develop and demonstrate a practical, scalable therapeutic for treating GIN infections in humans. The importance of these aspects of human anthelmintics are too often neglected. To date, there has been no drug developed specifically for human GINs. All drugs used for human GIN treatment came from drugs developed for veterinary targets (64). Furthermore, the availability of current drugs for human MDA relies upon off-patent drugs that are donated (22). Anthelmintics for human GINs need to be effective, safe, stable, scalable, compatible with MDA, and low-cost.

Bt Cry proteins, one of which has anthelmintic activity against human GINs (31, 34), have many of the characteristics required. Bt spore-crystal lysates are massively produced around the world for agriculture (75% of the biopesticide market; (25)), are shelf stable, and low-cost. Bt Cry proteins have a superb track record of safety for more than six decades of use in agriculture (caterpillar/beetle) and vector (black fly, mosquito) control (28, 29, 65). Indeed, the specific receptor for Cry5B in nematodes is restricted to invertebrates (35). But the use of Cry proteins as insecticidal sprays and in >100 MHa of transgenic crops at sub-anthelmintic doses is a “far cry” from their use as purposely-eaten therapeutics and as a broad spectrum anthelmintic.

Here, for the first time we describe a specific Bt Cry protein form developed to deliver Cry proteins as an ingestible therapeutic to vertebrates. This new form, Cry5B IBaCC, presents the Cry5B crystal protein as a crystal contained within the cell wall “ghost” of a dead vegetative bacterium, or paraprobiotic. Cry5B IBaCC is highly efficacious against multiple parasitic nematodes, including both genera of blood-feeding human hookworms and one lumenal-feeding parasitic nematode, *H. polygyrus bakeri*. IBaCC and/or the crystals inside the bacterium are ingested by the parasites, causing them to be intoxicated by damage to their intestinal cells. Efficacy *in vivo* is excellent, with a near complete clearance of hookworms at 15 mg/kg. On a molar concentration scale, Cry5B IBaCC is 175X more effective at clearing hookworms than albendazole (66).

Cry5B IBaCC production was successfully transferred to an industrial CMO and scaled up to 350 liters. Cry5B IBaCC is also compatible with simple fit-for-purpose formulations, such as sodium bicarbonate (*e.g*., Alka-Seltzer™) and can be dried down while retaining full activity. Importantly, this new form of Cry5B, IBaCC, appeared safe in a maximal multi-dose acute toxicology study based on histopathology and blood chemistry workups. These findings confirm the lack of Cry5B toxicity reported in all previous rodent, livestock, and companion animal studies (30–32, 34, 45). Our current focus is optimizing the Cry5B production strain for increased Cry5B yields, scale up, and GLP toxicology studies prior to initiating first-in-human clinical trials.

By killing the Cry5B-containing vegetative bacteria, but still using the whole fermentation, a host of issues associated with live bacteria or protein purification are obviated. Because the product is taken straight out of the fermenter, briefly incubated with essential oil, washed, and then ready for use, the process is simple and inexpensive. It could even be carried out locally in GIN endemic countries. Because the bacterium is killed, there are fewer issues with 1) degradation of the product over time as the bacterium dies, 2) regulatory issues associated with a product that is changing over time such as for live a bacteria as viability decays on the shelf, 3) selection of resistance with live bacteria replicating in the environment or in the GI tract, 4) release of live recombinant bacteria into the environment, and 5) any potential toxicity associated with enterotoxins associated with *B. cereus* family of bacteria. These properties should make IBaCC readily acceptable to drug and environmental regulatory agencies.

These studies validate IBaCC as a powerful, practical, safe, and deployable anthelmintic compatible with MDA that not only can safely and effectively deliver Cry5B but also any other anthelmintic Bt Cry protein for anti-GIN therapy. IBaCC uniquely and practically harnesses the safety, massive scalability, history, and power of Bt and Bt Cry proteins against one of the most prevalent and intractable disease of the poorest populations on earth. IBaCC promises new hope for a new arsenal of anthelmintics against the most common parasites of humans and animals.

## MATERIALS AND METHODS

### Nematodes

#### Medium and Reagents

Reagents for hookworm culture medium (HCM): RPMI 1640, fetal bovine serum (FBS), penicillin-streptomycin and fungizone antimycotic were all purchased from Gibco, U.S.A. Dexamethasone 21-phosphate disodium salt (DEX) (Cat# D1159-5G) and cimetidine (Cat# C4522-5G) were purchased from Sigma-Aldrich, USA. Cimetidine was prepared and dosed as described (36).

##### Caenorhabditis elegans

*Caenorhabditis elegans* was maintained using standard techniques (37). The following strains were used in this study: N2 Bristol (wild-type) and *glp-4(bn2)*. For images taken in Figure 3 (growth assay), hatched N2 L1 worms were incubated for 3 days at 25° C using a standard L1 growth assay (38, 39) in 48 well plates containing an *E. coli* food source and with treatments as indicated. Assayed worms were stilled with sodium azide, washed, and arranged for imaging in spot plates. Images were taken with a dissecting microscope fitted with a camera. The bioactivity of Cry5B in freeze-dried and irradiated freeze-dried SCLs (Fig. 1) was confirmed against *C. elegans* by a mortality assay for 48 hours at 25°C as described (39–41). For Fig. 3 lethality study, assays were carried out with *glp-4(bn2)* hermaphrodites incubated at 25°C for 6 days. Data represent the average and standard error from three independent experiments with approximately 60 worms per experiment (180 total), except for the IBa control in Fig. 3A (two independent experiments).

#### Animals and Parasites

*Ancylostoma ceylanicum* and *Necator americanus* life cycles were maintained as previously published (31). Three-to four-week-old male Golden Syrian hamsters (HsdHan:AURA) were purchased from Envigo (U.S.A) and were infected at approximately 4–5 weeks of age with either ~ 150 *A. ceylanicum* third-stage infectious larvae (L3i) orally or ~400 *N. americanus* L3i subcutaneously. Hamsters were provided with food and water *ad libitum*. The *Heligmosomoides polygyrus bakeri* life cycle was maintained at the United States Department of Agriculture (USDA) as described (42). Infectious staged larvae were shipped to University of Massachusetts Medical School. All animal experiments were carried out under protocols approved by the University of Massachusetts Medical School. All housing and care of laboratory animals used in this study conform to the NIH Guide for the Care and Use of Laboratory Animals in Research (see 18-F22) and all requirements and all regulations issued by the USDA, including regulations implementing the Animal Welfare Act (P.L. 89-544) as amended (see 18-F23).

#### *In vitro* assays with parasites

Egg-to-larva assays were carried out as described ((43); manuscript in preparation). Adult hookworm *in vitro* assays were carried out essentially as described (32, 44). Briefly, for *A. ceylanicum*, three adult hookworms per well were placed in 500 μL hookworm medium in a 24 well format with the indicated treatment using four wells/condition and then set up three independent times. *N. americanus* parasites were similarly tested but with only three wells per condition, because the number of adult parasites per hamster is more limited. For all conditions, there were roughly the same number of male and female worms. Hookworm adults were scored on a 0-3 scale (0 non-motile even when touched; 1 non-motile unless touched; 2 slowly motile; 3 fully motile) as described (32, 44).

Rhodamine-labeled IBaCC (rhod-IBaCC): IBaCC was resuspended in phosphate buffer (0.1 M, pH 7) at a concentration of 2 mg Cry5B/mL. Rhodamine isothiocyanate (RITC) was dissolved at 5 mg/mL in dimethylsulfoxide. RITC solution (60 μL) was added to a 1 mL suspension of IBaCC and the sample was incubated with constant mixing in the dark, at room temperature for 18 hours Tris buffer (60 μL, 1 M, pH 8) was added and the reaction mixture was stirred for additional 15 minutes to quench free RITC. The sample was centrifuged to collect rhod-IBaCC and the pellet was washed with water to remove physisorbed dye. IBaCC and rhod-IBaCC were evaluated for particle uptake in *A*. *ceylanicum* at a concentration of 15 μg Cry5B/mL using three adult hookworms per well in a 24-well format. Worms were evaluated by fluorescence microscopy for particle uptake at 1.5, 4 and 24 hours.

#### *In vivo* studies

The *A. ceylanicum* and *N. americanus in vivo* experiments were carried out as described (31–33, 44). Briefly, fecal egg counts for *A. ceylanicum* were taken day 17-18 post-inoculation to establish groups with roughly equal infectivity and then treated with a single dose gavage on day 18 post-inoculation. Fecal egg counts were taken again days 22-23 post-inoculation, and parasite burdens in the small and large intestine were taken on day 23 post-inoculation. Fecal egg counts for *N. americanus* were taken on days 55-56 post-inoculation to establish groups with roughly equal infectivity and then treated with a single dose gavage on day 56 post-inoculation. Fecal egg counts were taken again on days 60-61 post-inoculation, and parasite burdens in the small and large intestine were taken on day 61 post-inoculation. For all *in vivo* experiments except where sodium bicarbonate was used (Fig. 7A; all of Fig. 8; Fig. S1), cimetidine was prepared and given *per os* 15 minutes ahead of Cry5B administration as previously described (31). For experiments with sodium bicarbonate, Cry5B IBaCC was given *per os* in 200 μL 0.1 M sodium bicarbonate. Freeze-dried IBaCC was prepared as for SCL (described below). Experiments using *H. polygyrus bakeri*, experiments were carried out as described (31). Briefly, fecal egg counts were taken on days 14-15 post-inoculation to establish groups with roughly equal infectivity and then treated with a single dose gavage day on 15 post-inoculation. Fecal egg counts were taken again on days 19-20 post-inoculation, and parasite burdens in the small and large intestine were taken on day 20 post-inoculation.

### Bacteria

#### Spore and spore crystal lysates

For the experiments in Figure 1, Bt subspecies *kurstaki* HD1-4D8 and HD1-4D9 were ordered through the *Bacillus* Genetic Stock Center. Both crystal-deficient Bt strains were transformed with a plasmid containing the Cry5B gene (40). Spore lysates (SLs; HD1 Cry deficient strains) and spore-crystal lysates (SCLs; HD1 strains transformed with Cry5B plasmid) were prepared using standard methods (40, 45) and then stored at −80° C until use. For freeze drying, SL and SCL samples stored at −80°C for at least 12 hours were loaded into a FreeZone 1 Liter Benchtop Freeze Dry System (Labconco catalog number 7740020). The condenser was set to −60°C and the vacuum at 22 mTor. Irradiation of freeze-dried SL and SCL was accomplished with a cobalt-60 irradiation source at the University of Massachusetts Lowell Radiation Laboratory. Irradiation doses of 5, 10, 15, 20, 25, 30, and 60 kGy were initially tested, and 15kGy was chosen as the lowest radiation dose with a strong effect on spore viability. To determine the number of spores, 10 mg lyophilized powder was taken under sterile conditions before and after irradiation, transferred into 1 ml of sterile distilled water in microtubes, and homogenized by vortexing. 100 μl of the SL or SCL suspensions were removed from each sample and incubated at 80°C in a water bath for 20 minutes to kill any vegetative cells, and then diluted by a 10-fold dilution series. 100 μl of diluted samples (from 10^6^ to 10^9^) were spread on top of LB agar plates in triplicate with Rattler plating beads and incubated overnight at 30°C. Colonies on each plate were manually counted.

#### IBa and IBaCC strain construction and maintenance

The promoter region of *cry3A* (46) was fused to the coding region of *cry5B* and its downstream terminator via the N-terminal *ClaI* restriction site. This P*_cry3A_-cry5B* expression construct was subsequently cloned into pHT3101 (47). The resulting plasmid, pHY159, was electroporated into *B. thuringiensis* strain 407 Δ*spo0A::kan* (48). The entire *cry5B* insert was sequenced to confirm no mutations were included. Single colonies of 407 Δ*spo0A::kan* cells harboring pHY159 or pHT3101 empty vector control (EVC) were grown in nutrient-rich 3X LB with erythromycin (10ug/ml) at 30°C, shaking at 250rpm for 2-3 days. Outsourced cultures at a biomanufacturing facility were similarly grown in a fermenter with constant monitoring and adjustments of pH and oxygen levels, at 30°C with 150 rpm agitation for 48 hours. In both cases, cells were harvested by centrifugation and resuspended to 10% initial volumes in ice-cold water. 10X concentrations of harvested cultures were inactivated with the addition of food grade monoterpene or essential oil at 0.1% final concentration and incubated with gentle agitation at room temperature for 15 minutes. Inactivated cultures were then centrifuged, washed with water, and resuspended in saline at 10% initial volume. For all experiments except Fig. 7B, residual terpenes were extracted with corn oil (20% final volume) with gentle agitation at room temperature for 2 hours. We have not found any impact on *in vivo* efficacy with or without this step. IBaCC was recovered after centrifugation and three washes with ice-cold water. 10X concentrated samples were removed for several analyses, including SDS-PAGE (for Cry5B content and quantification), cell density, cell viability, and nematode-killing assays.

For experiments in Table S1, Bt 407 Δ*spo0A::kan* cells transferred with pHY159 were grown in Luria broth overnight at 30°C in 10 μg/mL erythromycin to saturation. The next day, 100 μL of terpenes (49) were added to 100 μL of overnight culture to a final volume of 1 mL in Luria broth plus erythromycin. Tubes were shaken overnight at 30°C. The next day, the cells were pelleted by centrifugation, washed with sterile water three times, and resuspended in 1 mL of sterile water. From each sample, 25 μL was plated onto a Luria broth plate and incubated at 30°C overnight. Growth or lack of growth was noted the following day.

#### Histopathology and Blood Chemistry

Organs from euthanized male and female animals were fixed in 10% neutral buffered formalin, processed and embedded in paraffin, sectioned at 5 μm, and stained with hematoxylin and eosin at Tufts University, Cummings School of Veterinary Medicine, Core Histology Laboratory (North Grafton, MA, USA). Hematoxylin stains nuclei and structures rich in nucleic acids gray to blue to dark purple. Eosin stains protein-rich regions of cells and tissues various shades of pink. All sections were blindly examined by a board-certified veterinary pathologist (GB). The organs sampled and (# sections) examined per animal included: stomach (2), small intestine (4), pancreas (1), large intestine (2–3), kidney (2), adrenal (1), liver (3), spleen (1), mesentery (1), brain (4–6), lungs (3–4), heart (entire), thymus (1–2), cortical bone (multiple), bone marrow (multiple), growth plate (multiple). All tissues were examined for microscopic disease and lesions as follows: Presence of nematodes/eggs; Cellular immune or inflammatory infiltrates; Cellular degeneration, apoptosis, necrosis; Lesion severity (none, minimal, moderate, severe); Lesion location (anatomic site); and incidental findings. After examination without knowledge of the groups, the study key was provided. In all animals and in all groups (males, females, IBaCC, and water), the following tissues were considered within normal limits by light microscopy: Lymphoid or hematopoietic tissues (thymus, spleen, Peyer’s Patches, bone marrow); Central nervous system (brain); Endocrine (adrenal); Skeletal (bone; growth plates); Cardiopulmonary (heart, lungs), upper digestive tract (Squamous portion of the stomach, liver, or pancreas), and lower digestive tract (large intestine). The small intestine of all animals in all treatment groups contained minimal to mild, multifocal, plasmocytic to lymphoplasmocytic and rarely eosinophilic infiltrates within the lamina propria. The infiltrates were interpreted as normal resident mucosal immune cells. There was no evidence of significant inflammation, degeneration, necrosis, fibrosis, or other toxicities in these tissues. Multifocal, minimal to mild, mineralized foci were noted in the glandular epithelium of the stomach and the kidneys (renal tubules and collecting ducts) of all males and females in both the IBaCC and water groups. The cause and clinical significance of the mineralization is uncertain. Regardless, mineralization did not appear to be a specific adverse effect attributable to IBaCC administration. Further studies are needed to determine the pathogenesis and clinical relevance of this lesion. The glandular stomach of females and males in both treatment groups contained particulates in the superficial mucus, of uncertain cause or significance. In a fraction of animals in all treatment groups (1/4 IBaCC treated females; 2/3 water-treated females; 3/3 IBaCC males and 2/4 water-treated males), minimal to mild lymphocytic serositis with reactive mesothelial cell hypertrophy was observed. The cause and clinical significance of this lesion was not apparent, but the lesion does not appear to be a specific adverse effect attributable to IBaCC treatment.

Immediately after euthanasia, hamsters were exsanguinated by cardiac puncture and blood collected into SAFE-T-FILL Capillary Blood Collection Tubes–Serum (RAM Scientific). Blood was allowed to clot for at least 30 min at room temperature before centrifugation. The collected serum was stored in −80°C until further use. Serum biochemistry profiles were performed on a COBAS c501 chemistry analyzer (Roche Diagnostics, Indianapolis, IN, USA) using standard protocols.

### 2.5. Statistical analyses

Prism v. 7 was used for all graphs and two group comparisons. Multigroup comparisons were carried out with SPSS v. 25. For all comparisons including just two groups except serum biochemistry (see Table S3), a one-tailed student’s t-test was used with the assumption that treatment reduced parasite burdens and fecal egg counts. For all comparisons involving two groups relative to a control group, one-way analysis of variance (ANOVA) with a one-tailed Dunnett’s post-test was used.

## ACKNOWLEDGEMENTS

This project was supported by (1) the National Institutes of Health/National Institute of Allergy and Infectious Diseases grants R01AI056189 and R01AI50866 to R.V.A., and (2) Agriculture and Food Research Initiative Competitive Grant no. 2015–11323 from the USDA National Institute of Food and Agriculture to R.V.A. We thank Ms Linda Wrijil and Ms Sarah Ducat for the excellent histology services at Tufts University’s Cummings School of Veterinary Medicine.

---

**Figure S1.**
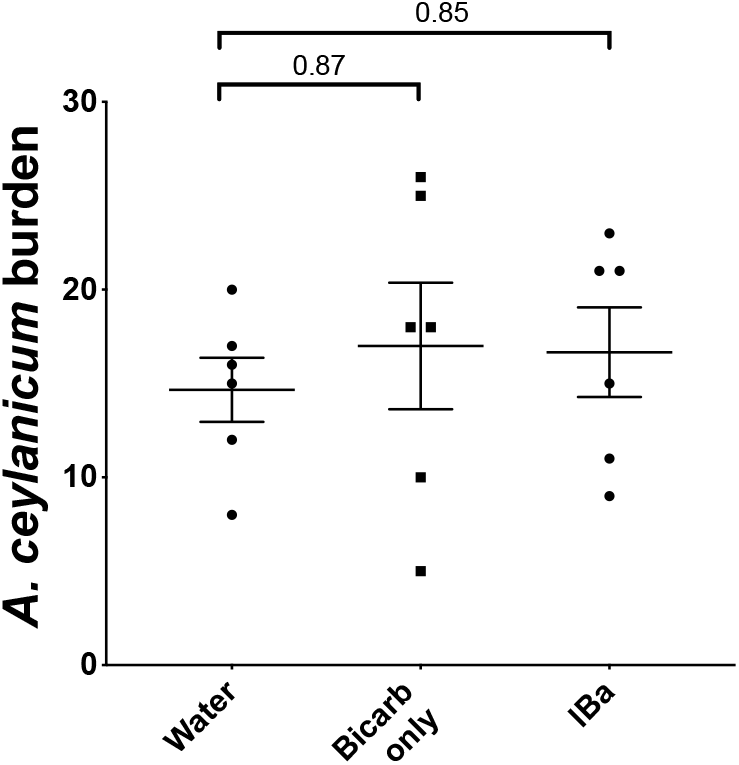
Negative control studies for hookworm burdens. Neither sodium bicarbonate alone nor IBa significantly impact hookworm burdens in hamsters.

**Figure S2.**
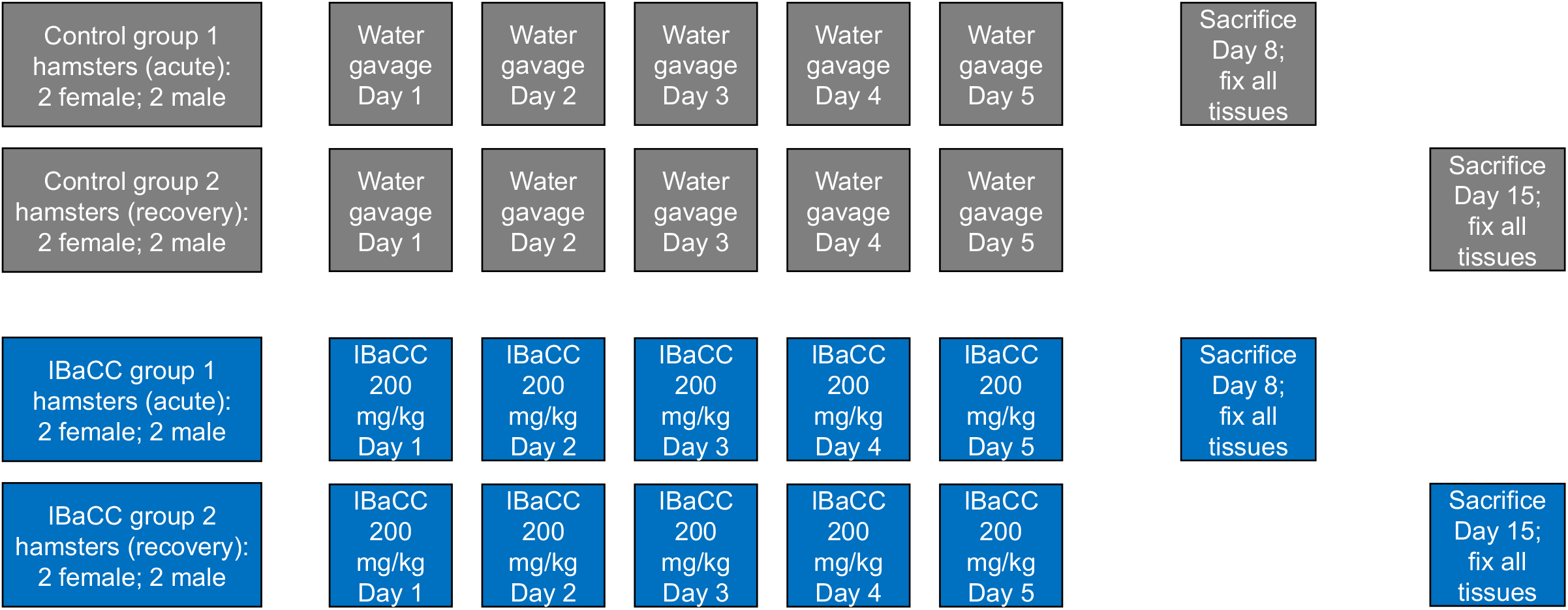
Experimental design for preliminary safety study. Volume of gavage was the same in all groups.

**Table S1.**
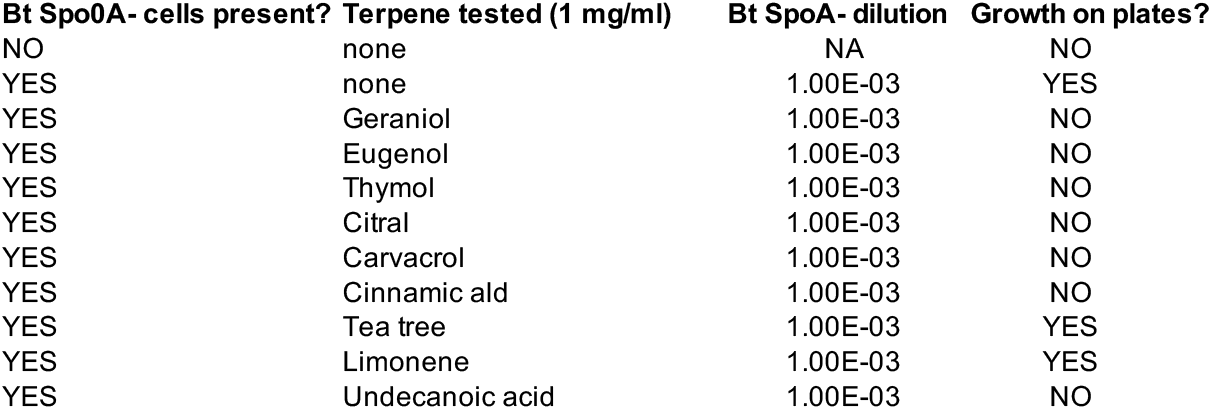
Qualitative survey of essential oils tested against BaCC cells.

**Table S2.**
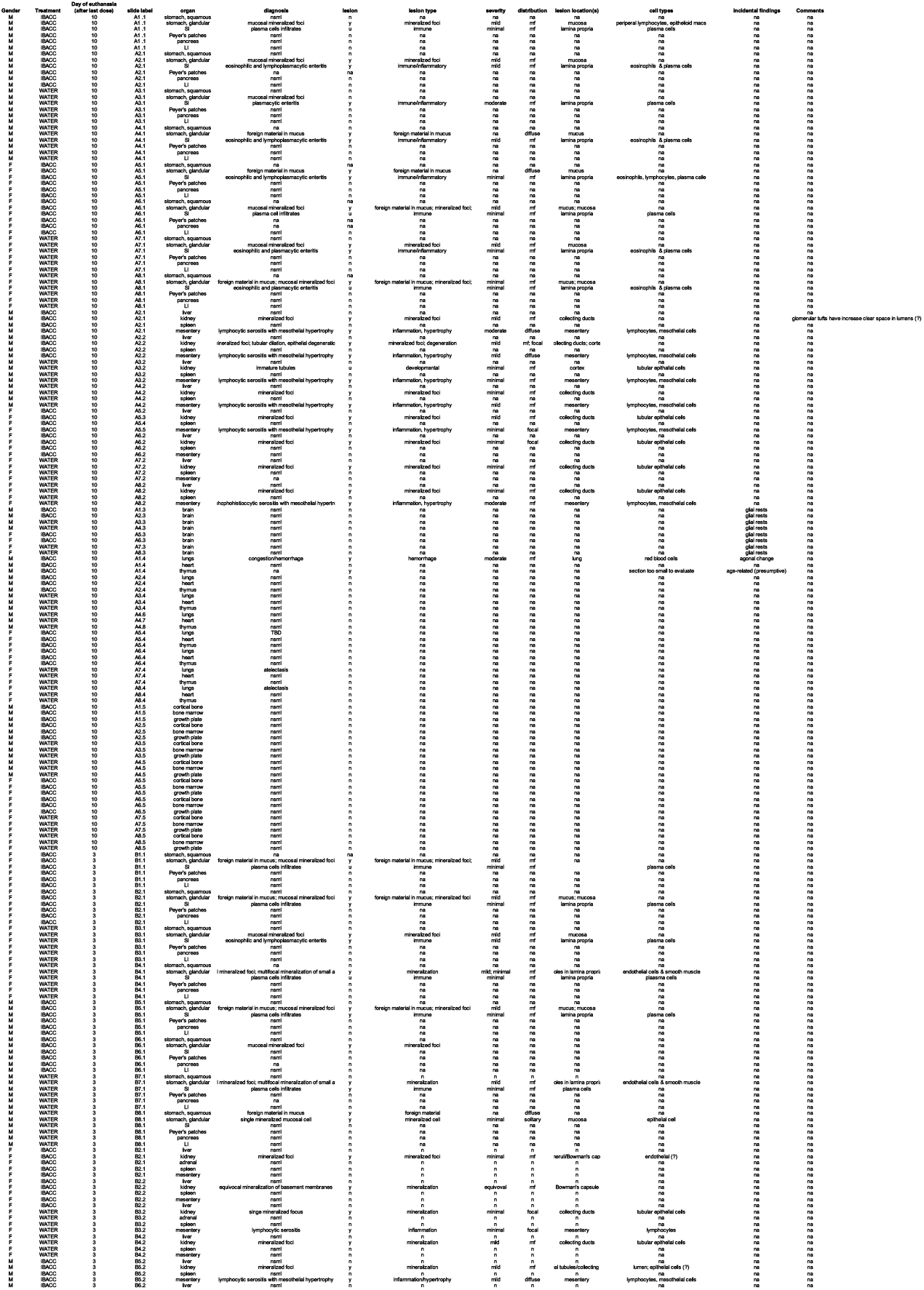
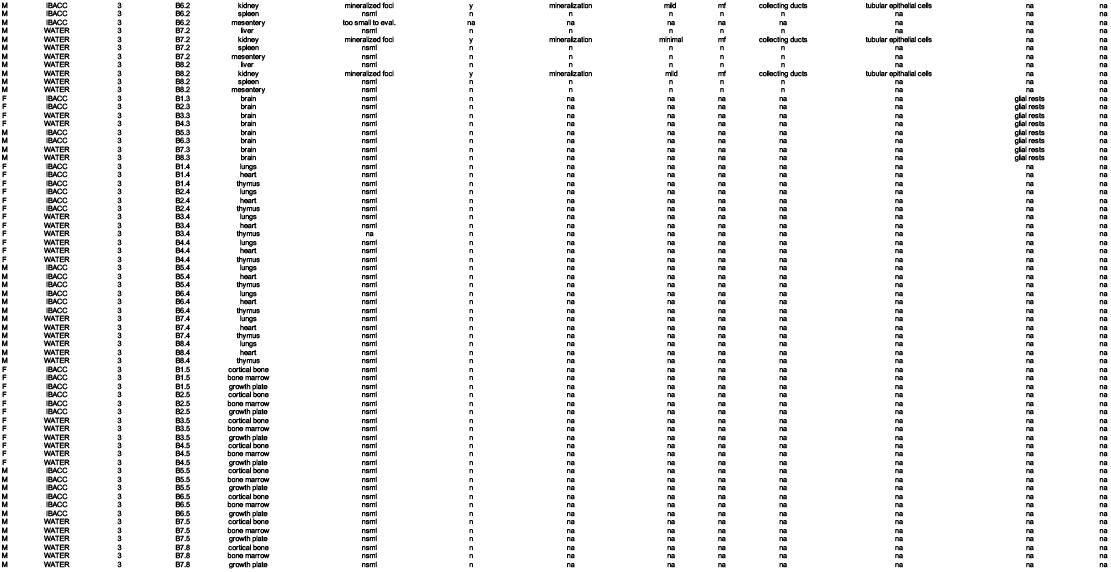
Histopathology results. nsml=No Significant Microscopic Lesions; na=that particular tissue was not available or seen for that animal.

**Table S3.**
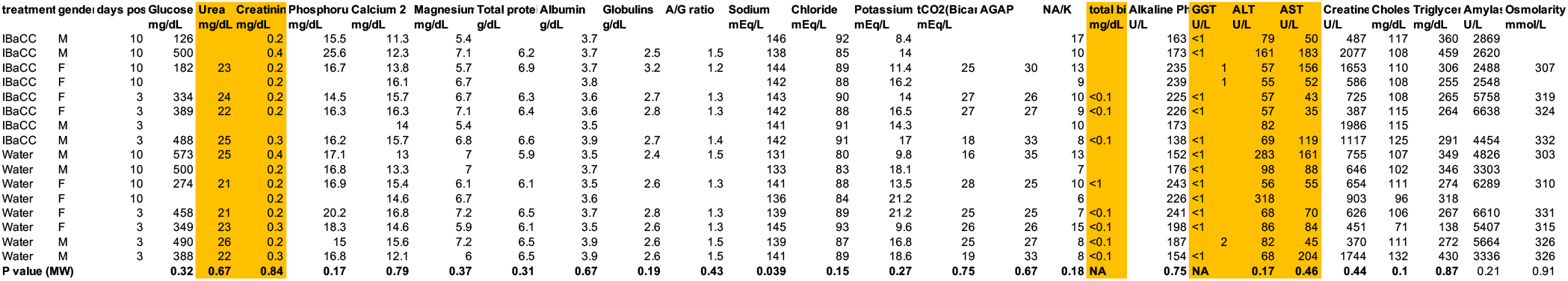
Serum biochemistry values for all animals in histopathology study. Missing values are due to limited amount of serum available for particular animals. P values are two-tailed Mann-Whitney (MW; http://www.statskingdom.com/170median_mann_whitney.html) comparing all Cry5B IBaCC-treated hamsters versus all Water-treated hamsters for any given parameter. Further breakdown (based on gender, days post last gavage or days post) are not possible due to limited number of animals per group. Only average sodium levels were statistically different (slightly elevated 142 mEq/L in IBaCC vs 138 mEq/L in Water). Columns in color are relevant for determining any potential hepatic and renal injury. NA = no statistical comparison possible due to nature readout but clearly no difference between IBaCC and Water groups is seen.

## REFERENCES

1. Schulz JD, Moser W, Hürlimann E, Keiser J. 2018. Preventive Chemotherapy in the Fight against Soil-Transmitted Helminthiasis: Achievements and Limitations. Trends Parasitol 34:590–602.

2. Bethony J, Brooker S, Albonico M, Geiger SM, Loukas A, Diemert D, Hotez PJ. 2006. Soil-transmitted helminth infections: ascariasis, trichuriasis, and hookworm. Lancet 367:1521–1532.

3. Ojha SC, Jaide C, Jinawath N, Rotjanapan P, Baral P. 2014. Geohelminths: public health significance. J Infect Dev Ctries 8:5–16.

4. Jourdan PM, Lamberton PHL, Fenwick A, Addiss DG. 2018. Soil-transmitted helminth infections. Lancet 391:252–265.

5. Hotez P. 2008. Hookworm and poverty. Ann N Y Acad Sci 1136:38–44.

6. Passerini L, Casey GJ, Biggs BA, Cong DT, Phu LB, Phuc TQ, Carone M, Montresor A. 2012. Increased birth weight associated with regular pre-pregnancy deworming and weekly iron-folic acid supplementation for Vietnamese women. PLoS Negl Trop Dis 6:e1608.

7. Pullan RL, Smith JL, Jasrasaria R, Brooker SJ. 2014. Global numbers of infection and disease burden of soil transmitted helminth infections in 2010. Parasit Vectors 7:37.

8. Keiser J, Utzinger J. 2010. The drugs we have and the drugs we need against major helminth infections. Adv Parasitol 73:197–230.

9. Keiser J, Utzinger J. 2008. Efficacy of current drugs against soil-transmitted helminth infections: systematic review and meta-analysis. JAMA 299:1937–1948.

10. Kaplan RM. 2004. Drug resistance in nematodes of veterinary importance: a status report. Trends Parasitol 20:477–481.

11. Humphries D, Mosites E, Otchere J, Twum WA, Woo L, Jones-Sanpei H, Harrison LM, Bungiro RD, Benham-Pyle B, Bimi L, Edoh D, Bosompem K, Wilson M, Cappello M. 2011. Epidemiology of hookworm infection in Kintampo North Municipality, Ghana: patterns of malaria coinfection, anemia, and albendazole treatment failure. Am J Trop Med Hyg 84:792–800.

12. Humphries D, Simms BT, Davey D, Otchere J, Quagraine J, Terryah S, Newton S, Berg E, Harrison LM, Boakye D, Wilson M, Cappello M. 2013. Hookworm infection among school age children in Kintampo north municipality, Ghana: nutritional risk factors and response to albendazole treatment. Am J Trop Med Hyg 89:540–548.

13. Humphries D, Nguyen S, Kumar S, Quagraine JE, Otchere J, Harrison LM, Wilson M, Cappello M. 2017. Effectiveness of Albendazole for Hookworm Varies Widely by Community and Correlates with Nutritional Factors: A Cross-Sectional Study of School-Age Children in Ghana. Am J Trop Med Hyg 96:347–354.

14. Stothard JR, French MD, Khamis IS, Basáñez M-G, Rollinson D. 2009. The epidemiology and control of urinary schistosomiasis and soil-transmitted helminthiasis in schoolchildren on Unguja Island, Zanzibar. Trans R Soc Trop Med Hyg 103:1031–1044.

15. Stothard JR, Rollinson D, Imison E, Khamis IS. 2009. A spot-check of the efficacies of albendazole or levamisole, against soil-transmitted helminthiases in young Ungujan children, reveals low frequencies of cure. Ann Trop Med Parasitol 103:357–360.

16. Soukhathammavong PA, Sayasone S, Phongluxa K, Xayaseng V, Utzinger J, Vounatsou P, Hatz C, Akkhavong K, Keiser J, Odermatt P. 2012. Low efficacy of single-dose albendazole and mebendazole against hookworm and effect on concomitant helminth infection in Lao PDR. PLoS Negl Trop Dis 6:e1417.

17. Edelduok EG, Eke FN, Evelyn NE, Atama CI, Eyo JE. 2013. Efficacy of a single dose albendazole chemotherapy on human intestinal helminthiasis among school children in selected rural tropical communities. Annals of Tropical Medicine and Public Health 6:413.

18. Krücken J, Fraundorfer K, Mugisha JC, Ramünke S, Sifft KC, Geus D, Habarugira F, Ndoli J, Sendegeya A, Mukampunga C, Bayingana C, Aebischer T, Demeler J, Gahutu JB, Mockenhaupt FP, von Samson-Himmelstjerna G. 2017. Reduced efficacy of albendazole against Ascaris lumbricoides in Rwandan schoolchildren. Int J Parasitol Drugs Drug Resist 7:262–271.

19. Diawara A, Halpenny CM, Churcher TS, Mwandawiro C, Kihara J, Kaplan RM, Streit TG, Idaghdour Y, Scott ME, Basáñez M-G, Prichard RK. 2013. Association between response to albendazole treatment and β-tubulin genotype frequencies in soil-transmitted helminths. PLoS Negl Trop Dis 7:e2247.

20. Zuccherato LW, Furtado LF, Medeiros C da S, Pinheiro C da S, Rabelo ÉM. 2018. PCR-RFLP screening of polymorphisms associated with benzimidazole resistance in Necator americanus and Ascaris lumbricoides from different geographical regions in Brazil. PLoS Negl Trop Dis 12:e0006766.

21. DiMasi JA, Grabowski HG, Hansen RW. 2016. Innovation in the pharmaceutical industry: New estimates of R&D costs. J Health Econ 47:20–33.

22. Lin WM, Addiss DG. 2018. Sustainable access to deworming drugs in a changing landscape. Lancet Infect Dis 18:e395–e398.

23. Dayan AD. 2003. Albendazole, mebendazole and praziquantel. Review of non-clinical toxicity and pharmacokinetics. Acta Trop 86:141–159.

24. 2019. MebendazoleLiverTox: Clinical and Research Information on Drug-Induced Liver Injury. National Institute of Diabetes and Digestive and Kidney Diseases, Bethesda (MD).

25. Karthickumar P, Balasubramanian P. 2017. Biofertilizers and Biopesticides: A Holistic Approach for Sustainable Agriculture, p. 269–298. *In* Sustainable Utilization of Natural Resources. CRC Press.

26. Griffitts JS, Aroian RV. 2005. Many roads to resistance: how invertebrates adapt to Bt toxins. Bioessays 27:614–624.

27. Tabashnik BE, Carrière Y. 2017. Surge in insect resistance to transgenic crops and prospects for sustainability. Nat Biotechnol 35:926–935.

28. Betz FS, Hammond BG, Fuchs RL. 2000. Safety and advantages of Bacillus thuringiensis-protected plants to control insect pests. Regul Toxicol Pharmacol 32:156–173.

29. Koch MS, Ward JM, Levine SL, Baum JA, Vicini JL, Hammond BG. 2015. The food and environmental safety of Bt crops. Front Plant Sci 6:283.

30. Cappello M, Bungiro RD, Harrison LM, Bischof LJ, Griffitts JS, Barrows BD, Aroian RV. 2006. A purified Bacillus thuringiensis crystal protein with therapeutic activity against the hookworm parasite Ancylostoma ceylanicum. Proc Natl Acad Sci U S A 103:15154–15159.

31. Hu Y, Nguyen T-T, Lee ACY, Urban JF Jr, Miller MM, Zhan B, Koch DJ, Noon JB, Abraham A, Fujiwara RT, Bowman DD, Ostroff GR, Aroian RV. 2018. Bacillus thuringiensis Cry5B protein as a new pan-hookworm cure. Int J Parasitol Drugs Drug Resist 8:287–294.

32. Hu Y, Zhan B, Keegan B, Yiu YY, Miller MM, Jones K, Aroian RV. 2012. Mechanistic and single-dose in vivo therapeutic studies of Cry5B anthelmintic action against hookworms. PLoS Negl Trop Dis 6:e1900.

33. Hu Y, Miller MM, Derman AI, Ellis BL, Monnerat RG, Pogliano J, Aroian RV. 2013. Bacillus subtilis strain engineered for treatment of soil-transmitted helminth diseases. Appl Environ Microbiol 79:5527–5532.

34. Urban JF Jr, Hu Y, Miller MM, Scheib U, Yiu YY, Aroian RV. 2013. Bacillus thuringiensis-derived Cry5B has potent anthelmintic activity against Ascaris suum. PLoS Negl Trop Dis 7:e2263.

35. Griffitts JS, Haslam SM, Yang T, Garczynski SF, Mulloy B, Morris H, Cremer PS, Dell A, Adang MJ, Aroian RV. 2005. Glycolipids as receptors for Bacillus thuringiensis crystal toxin. Science 307:922–925.

36. Stepek G, Lowe AE, Buttle DJ, Duce IR, Behnke JM. 2007. Anthelmintic action of plant cysteine proteinases against the rodent stomach nematode, Protospirura muricola, in vitro and in vivo. Parasitology 134:103–112.

37. Brenner S. 1974. The genetics of Caenorhabditis elegans. Genetics 77:71–94.

38. Wei J-Z, Hale K, Carta L, Platzer E, Wong C, Fang S-C, Aroian RV. 2003. Bacillus thuringiensis crystal proteins that target nematodes. Proc Natl Acad Sci U S A 100:2760–2765.

39. Bischof LJ, Huffman DL, Aroian RV. 2006. Assays for toxicity studies in C. elegans with Bt crystal proteins. Methods Mol Biol 351:139–154.

40. Marroquin LD, Elyassnia D, Griffitts JS, Feitelson JS, Aroian RV. 2000. Bacillus thuringiensis (Bt) toxin susceptibility and isolation of resistance mutants in the nematode Caenorhabditis elegans. Genetics 155:1693–1699.

41. Hu Y, Platzer EG, Bellier A, Aroian RV. 2010. Discovery of a highly synergistic anthelmintic combination that shows mutual hypersusceptibility. Proc Natl Acad Sci U S A 107:5955–5960.

42. Camberis M, Le Gros G, Urban J Jr. 2003. Animal model of Nippostrongylus brasiliensis and Heligmosomoides polygyrus. Curr Protoc Immunol Chapter 19:Unit 19.12.

43. Hu Y, Miller M, Zhang B, Nguyen T-T, Nielsen MK, Aroian RV. 2018. In vivo and in vitro studies of Cry5B and nicotinic acetylcholine receptor agonist anthelmintics reveal a powerful and unique combination therapy against intestinal nematode parasites. PLoS Negl Trop Dis 12:e0006506.

44. Hu Y, Ellis BL, Yiu YY, Miller MM, Urban JF, Shi LZ, Aroian RV. 2013. An extensive comparison of the effect of anthelmintic classes on diverse nematodes. PLoS One 8:e70702.

45. Hu Y, Georghiou SB, Kelleher AJ, Aroian RV. 2010. Bacillus thuringiensis Cry5B protein is highly efficacious as a single-dose therapy against an intestinal roundworm infection in mice. PLoS Negl Trop Dis 4:e614.

46. Agaisse H, Lereclus D. 1994. Structural and functional analysis of the promoter region involved in full expression of the cryIIIA toxin gene of Bacillus thuringiensis. Mol Microbiol 13:97–107.

47. Lereclus D, Arantès O, Chaufaux J, Lecadet M. 1989. Transformation and expression of a cloned delta-endotoxin gene in Bacillus thuringiensis. FEMS Microbiol Lett 51:211–217.

48. Lereclus D, Agaisse H, Gominet M, Chaufaux J. 1995. Overproduction of Encapsulated Insecticidal Crystal Proteins in a Bacillus thuringiensis spoOA Mutant. Nat Biotechnol 13:67–71.

49. Mirza Z., Soto E.R., Hu Y., Nguyen T.T., Koch D., Aroian R.V., Ostroff G.R. 2020. Anthelmintic Activity of Yeast Particle-Encapsulated Terpenes. Molecules 25:2958.

50. Taverniti V, Guglielmetti S. 2011. The immunomodulatory properties of probiotic microorganisms beyond their viability (ghost probiotics: proposal of paraprobiotic concept). Genes Nutr 6:261–274.

51. Becker N. 2002. Sterilization of Bacillus thuringiensis israelensis products by gamma radiation. J Am Mosq Control Assoc 18:57–62.

52. Bevilacqua A, Corbo MR, Sinigaglia M. 2010. In vitro evaluation of the antimicrobial activity of eugenol, limonene, and citrus extract against bacteria and yeasts, representative of the spoiling microflora of fruit juices. J Food Prot 73:888–894.

53. Kang J, Liu L, Wu X, Sun Y, Liu Z. 2018. Effect of thyme essential oil against Bacillus cereus planktonic growth and biofilm formation. Appl Microbiol Biotechnol 102:10209–10218.

54. Gilling DH, Ravishankar S, Bright KR. 2019. Antimicrobial efficacy of plant essential oils and extracts against Escherichia coli. J Environ Sci Health A Tox Hazard Subst Environ Eng 54:608–616.

55. Soto E, Ostroff G. 2011. Use of□-Glucans for Drug Delivery Applications. Biology and Chemistry of Beta Glucan: Beta Glucans-Mechanisms of Action 1:48.

56. Wu C-C, Hu Y, Miller M, Aroian RV, Sailor MJ. 2015. Protection and Delivery of Anthelmintic Protein Cry5B to Nematodes Using Mesoporous Silicon Particles. ACS Nano 9:6158–6167.

57. Conlan JV, Khamlome B, Vongxay K, Elliot A, Pallant L, Sripa B, Blacksell SD, Fenwick S, Thompson RCA. 2012. Soil-transmitted helminthiasis in Laos: a community-wide cross-sectional study of humans and dogs in a mass drug administration environment. Am J Trop Med Hyg 86:624–634.

58. Schär F, Inpankaew T, Traub RJ, Khieu V, Dalsgaard A, Chimnoi W, Chhoun C, Sok D, Marti H, Muth S, Odermatt P. 2014. The prevalence and diversity of intestinal parasitic infections in humans and domestic animals in a rural Cambodian village. Parasitol Int 63:597–603.

59. Inpankaew T, Schär F, Dalsgaard A, Khieu V, Chimnoi W, Chhoun C, Sok D, Marti H, Muth S, Odermatt P, Traub RJ. 2014. High prevalence of Ancylostoma ceylanicum hookworm infections in humans, Cambodia, 2012. Emerg Infect Dis 20:976–982.

60. Bradbury RS, Hii SF, Harrington H, Speare R, Traub R. 2017. Ancylostoma ceylanicum Hookworm in the Solomon Islands. Emerg Infect Dis 23:252–257.

61. Garside P, Behnke JM. 1989. Ancylostoma ceylanicum in the hamster: observations on the host—parasite relationship during primary infection. Parasitology 98:283–289.

62. Behnke JM, Rose R, Garside P. 1993. Sensitivity to ivermectin and pyrantel of Ancylostoma ceylanicum and Necator americanus. Int J Parasitol 23:945–952.

63. Bansemir AD, Sukhdeo MV. 1994. The food resource of adult Heligmosomoides polygyrus in the small intestine. J Parasitol 80:24–28.

64. Elfawal, M. A., Savinov, S.N., and Aroian, R.V. 2019. Drug Screening for Discovery of Broad-spectrum Agents for Soil-transmitted Nematodes. Sci Rep in review.

65. Roh JY, Choi JY, Li MS, Jin BR, Je YH. 2007. Bacillus thuringiensis as a specific, safe, and effective tool for insect pest control. J Microbiol Biotechnol 17:547–559.

66. Tritten L, Silbereisen A, Keiser J. 2011. In vitro and in vivo efficacy of Monepantel (AAD 1566) against laboratory models of human intestinal nematode infections. PLoS Negl Trop Dis 5:e1457.

